# NEST Desktop - An educational application for neuroscience

**DOI:** 10.1101/2021.06.15.444791

**Authors:** Sebastian Spreizer, Johanna Senk, Stefan Rotter, Markus Diesmann, Benjamin Weyers

**Affiliations:** Faculty of Biology, University of Freiburg, 79104 Freiburg, Germany; Bernstein Center Freiburg, University of Freiburg, 79104 Freiburg, Germany; Institute of Neuroscience and Medicine (INM-6) and Institute for Advanced Simulation (IAS-6) and JARA-Institute Brain Structure-Function Relationships (INM-10), Jülich Research Centre, 52428 Jülich, Germany; Department of Computer Science, University of Trier, 54296 Trier, Germany; Department of Psychiatry, Psychotherapy and Psychosomatics, School of Medicine, RWTH Aachen University, 52074 Aachen, Germany; Department of Physics, Faculty 1, RWTH Aachen University, 52074 Aachen, Germany

## Abstract

Simulation software for spiking neuronal network models matured in the past decades regarding performance and flexibility. But the entry barrier remains high for students and early career scientists in computational neuroscience since these simulators typically require programming skills and a complex installation. Here, we describe an installation-free Graphical User Interface (GUI) running in the web browser, which is distinct from the simulation engine running anywhere, on the student’s laptop or on a supercomputer. This architecture provides robustness against technological changes in the software stack and simplifies deployment for self-education and for teachers. Our new open source tool, NEST Desktop, comprises graphical elements for creating and configuring network models, running simulations, and visualizing and analyzing the results. NEST Desktop allows students to explore important concepts in computational neuroscience without the need to learn a simulator control language before. Our experiences so far highlight that NEST Desktop helps advancing both quality and intensity of teaching in computational neuroscience in regular university courses. We view the availability of the tool on public resources like the European ICT infrastructure for neuroscience EBRAINS as a contribution to equal opportunities.

**Significance Statement:** The graphical user interface NEST Desktop makes neuronal network simulations accessible to non-programmers. It facilitates the interactive exploration of neuronal network models by integrating the whole workflow of wiring up the setup, simulating the neuronal dynamics, and analyzing the recorded activity data into a single tool. NEST Desktop effectively supports teaching the concepts and methods of computational neuroscience. Due to its installation-free web-based implementation, it is in particularly suitable for online courses.

## Introduction

Complementary to experiment and theory, simulations of computational models represent an essential research tool in neuroscience. Neuronal network models integrate available knowledge of the brain’s individual constituents and their complex interactions with the aim to simulate neuronal activity matching data observed experimentally (Tikidji-Hamburyan et al., 2017). Dedicated open-source software tools, partially with decades of ongoing development and maintenance (Brette et al., 2007), promote the reproducibility of simulations, reuse and extension of code, and efficient usage of hardware. Many of these tools rely on textual, general-purpose programming languages (Einevoll et al., 2019) primarily designed for routine use by specialized researchers. Computational neuroscience, however, is an interdisciplinary field and scientists without strong background in programming often struggle to get started with the concepts and usage of simulators. Often enough they shipwreck already due to the complex installation of the software. To lower the entry barrier for these tools, and to provide access for non-programmers, a number of simulation engines have been equipped with Graphical User Interfaces (GUIs) to easily control simulations or explore network activity, see Table 1 for an overview.

**Table 1:** History of GUI development in computational neurosciene. GUIs are ordered chronologically according to the estimated beginning of their development phase (mentioned in a paper or first commit in a public repository). Most GUIs are coupled with a simulation engine in a specific front end environment. If a GUI is independent of specific simulators, the respective entry is “none”. “none (NEST)” in the case of VisNEST means that the application has full operational function without the simulator NEST but it can be connected to it. For more information, the last column lists the corresponding publications or refers to the source code.

| Development | GUI | Simulator | Environment | Reference |
| --- | --- | --- | --- | --- |
| 1992 | GENESIS GUI | GENESIS | x11 | Bower and Beeman (2012) |
| 1993 | NEURON GUI | NEURON | x11 | Hines (1993); Hines and Carnevale (1997) |
| 1995 | SLIDE | NEST | x11 | Matyak (1996); Gewaltig et al. (1996) |
| 2007 | neuroConstruct | multiple | x11 | Gleeson et al. (2007) |
| 2008 | SNN3DViewer | none | x11 | Kasiński et al. (2009) |
| 2009 | Neuronvisio | NEURON | x11 (qt4) | Mattioni et al. (2012) |
| 2011 | nuSPIC | NEST | HTML | Vlachos et al. (2013) |
| 2012 | The Virtual Brain (TVB) | TVB | HTML | Sanzleon et al. (2013) |
| 2013 | N2A (Neurons to Algorithms) | multiple | x11 (qt5) | Rothganger et al. (2014) |
| 2013 | SpineCreator | PyNN | x11 (qt5) | Cope et al. (2017) |
| 2013 | VisNEST | none (NEST) | VR | Nowke et al. (2013) |
| 2014 | Neuronify | Neuronify | x11 (qt5) | Dragly et al. (2017) |
| 2014 | Open Source Brain (OBS) | PyNN | HTML | Gleeson et al. (2019) |
| 2015 | Nengo GUI | Nengo | HTML | Source code<sup>1</sup> |
| 2015 | ViSimpl | none | x11 (qt5) | Galindo et al. (2016) |
| 2016 | **NEST Desktop** | NEST | HTML | Source code<sup>2</sup> |
| 2016 | VIOLA | none | HTML | Senk et al. (2018) |
| 2016 | Visbrain | none | x11 (qt5) | Combrisson et al. (2019) |
| 2017 | NESTInstrumentationApp | NEST | HTML | Source code<sup>3</sup> |
| 2017 | NetPyNE UI | NetPyNE | HTML | Dura-Bernal et al. (2019) |
| 2017 | NEURON UI | NEURON | HTML | Source code<sup>4</sup> |
| 2018 | CellExplorer | none | x11 (qt5) | Petersen et al. (2020) |

With our present work, we focus on college and university students as a specific user group where significant programming skills cannot be assumed. We present a web-based software tool, which has been specifically developed to support education and training of basic computational neuroscience for individual learners and classroom teaching. In addition, it is suited for online courses. The main educational objective is to develop solid understanding of how numerical simulations can be employed as a meaningful research tool in neuroscience. The methodological question is how the anatomy, physiology, and biophysics of neuronal systems should be translated into specific algorithmic components of a numerical simulation. Our didactic strategy is to enable exciting hands-on experience and rewarding results without delay and without big effort. The concept appeals to common sense and scientific intuition. It makes students enjoy the lessons, and invites independent creative research. For a successful implementation, we seek a framework that fulfills the following requirements. First, the tool needs to offer functionality that enables students to create a neuronal network model visually and interactively. Thus, there is no need for programming at this stage and the focus lies more on the neuroscientific questions. Second, there is the need to inspect the simulation results, in the sense of a constructive approach in learning (Clark and Mayer, 2011; de Jong et al., 2013). For this purpose, a simulator needs to be loaded with the model and executed, and it should then offer an easy-to-understand presentation of the results. This can then be the basis of a new iteration with an adapted model. Third, the tool should offer the use of standard models, storing and loading previous models as well as the creation of reports. Finally, the tool needs a high level of usability, should be easy to install, and scale up to a classroom size number of users.

The tool presented in this work is called NEST Desktop and aims to convey the structural and dynamical behavior of biological neuronal networks by addressing the requirements listed above. Building on what students with a general background in neuroscience are already familiar with, virtual experiments are conducted in similar steps, and described with similar terminology as known from biological experiments. Designed around the concept of constructivistic experimenting and fast prototyping, NEST Desktop allows users to explore key aspects of neuronal network modeling via a GUI: network construction and parameterization, running simulations, data analysis, and documentation can all be performed by means of interactive visualization (Fig. 1). In the background, NEST Desktop uses the NEural Simulation Tool (NEST, Gewaltig and Diesmann, 2007) as a reference simulation engine. NEST focuses on small to large networks of spiking neurons and comprises a collection of simple and more biophysically detailed neuron models. NEST Desktop is installation-free and requires only a modern web browser.

**Figure 1:**
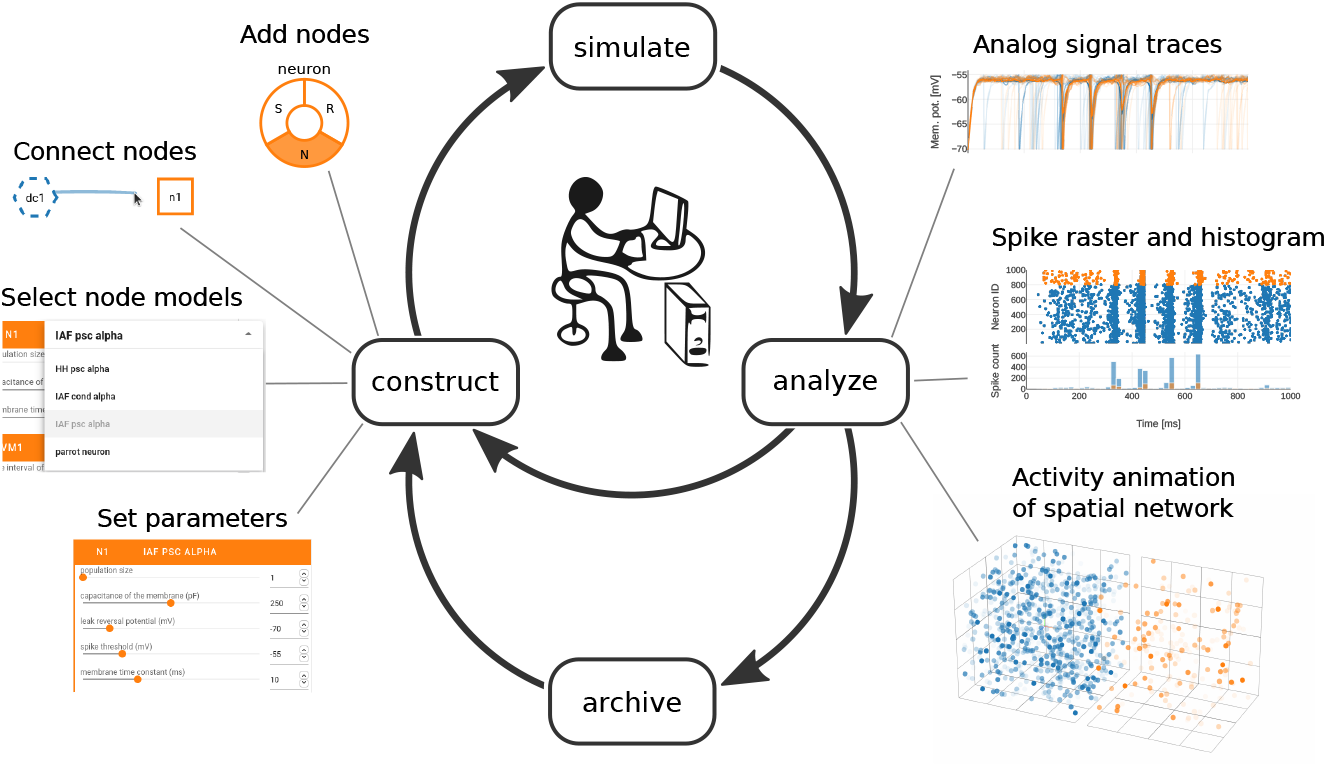
NEST Desktop workflow enables fast prototyping of neuronal network simulations. The GUI allows the user to control network construction, simulation, data analysis, and archiving via graphical elements. All these steps can be executed consecutively and repeatedly to explore model properties and resulting network dynamics. Constructing a network involves the selection and connection of nodes as well as their parameterization (‘construct’, left). After the simulation of a constructed network (‘simulate’, top), recorded analog signals and spiking activity can be assessed in various charts (‘analyze’, right). The user is also able to export and archive networks and results for documentation and later use (‘archive’, bottom).

The manuscript is structured as follows: The section Materials and Methods elucidates technical details of NEST Desktop. In Results, we describe the main components and functionality of NEST Desktop, and exemplify its usage with a use case about teaching in a classroom and supporting research. We have previously employed NEST Desktop in university courses and were able to make the experience that NEST Desktop successfully supports students to complete the course also in times of online courses due to the COVID-19 pandemic. The Discussion embeds the work in the state of research and reflects on general achievements, current limitations, and potential future developments.

Preliminary results have been published in abstract form (Spreizer, 2018; Spreizer et al., 2019, 2020).

## Materials and Methods

### Client-server architecture

NEST Desktop uses a client-server architecture (Fig. 2A,B): the client provides the GUI as front end and handles network construction, simulation-code generation, and analysis of activity data (Fig. 2B, purple row); the server runs the simulation engine NEST (Gewaltig and Diesmann, 2007) as back end and executes the simulation code (Fig. 2B, yellow row). This separation enables a lightweight and platform-independent implementation of NEST Desktop as a web application. The deployment of a NEST installation and an execution environment with sufficient computational resources is provided by the running infrastructure and is therefore outside of the user’s responsibility. However, this architecture requires the back end to remain stateless, which means that the data of the simulation (network model and simulation results) are stored only on the client side.

**Figure 2:**
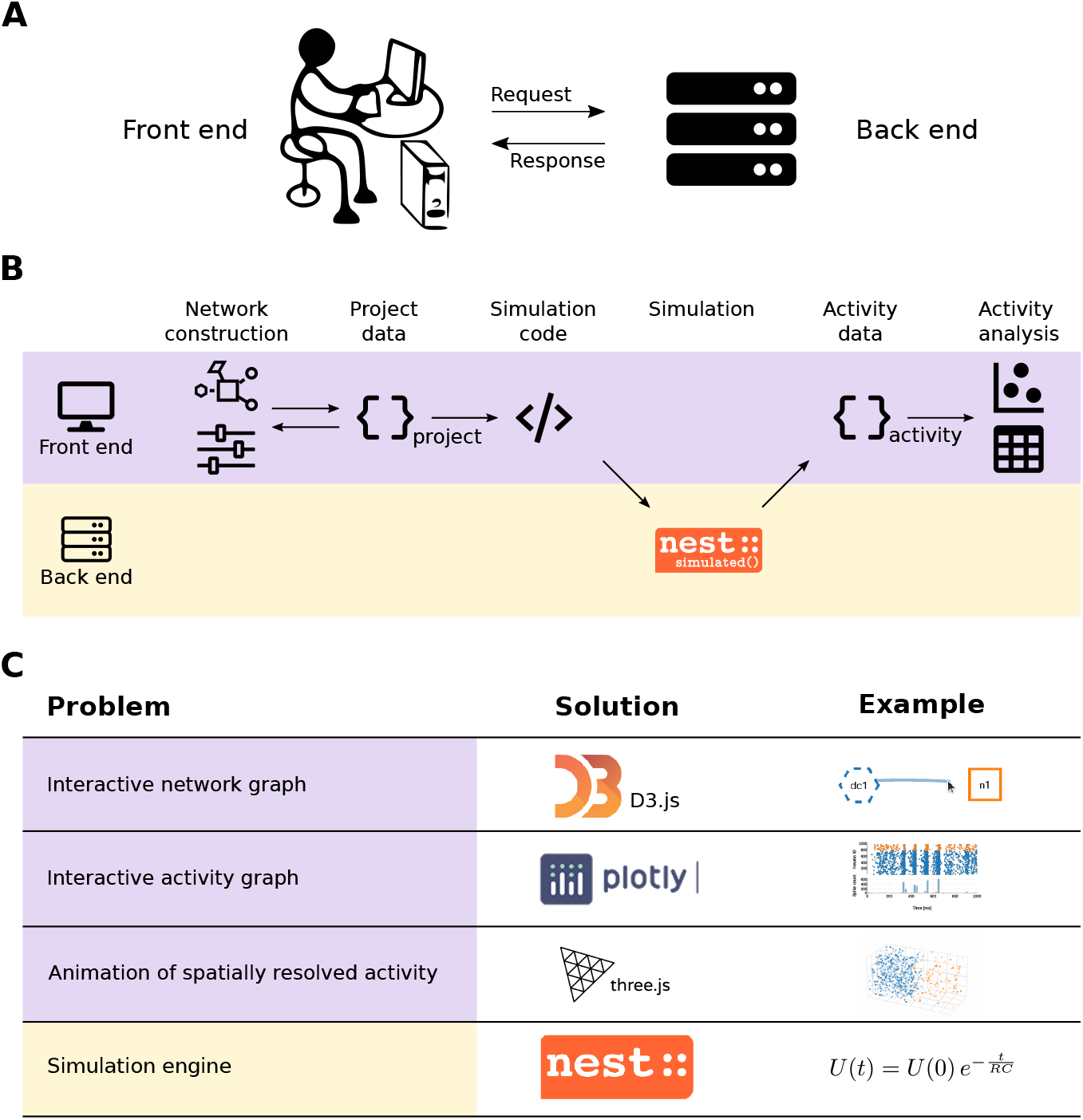
Workflow in client-server architecture and technical challenges. **(A)** Simplified relationship of the client (front end) and the server (back end). **(B)** The front end handles network construction and data analysis, whereas the back end executes simulations with NEST (Gewaltig and Diesmann, 2007). **(C)** Problem formulations regarding the software architecture (left column) together with the technical solutions implemented in NEST Desktop (center column) and associated examples (right column).

Standard data formats are used for communication to ensure compatibility of these front end and back end systems. NEST 3 (Hahne et al., 2021) offers ‘API server’ as a server-side wrapper of NEST Simulator in Python, which enables the communication of NEST Desktop as GUI on the client side (i.e., in the browser) and ‘NEST Simulator’ (or just ‘NEST’) as simulation engine on the server side. NEST Desktop and NEST Simulator use JSON for the server-to-browser (stateless) communication over standard HTTP as communication protocol. JSON is a language-independent data format which can be applied in JavaScript, enabling interactive web pages, and is interpretable by most programming languages including Python.

### Front end implementation

The GUI as a front end (Fig. 2, purple rows) makes use of modern web technologies (e.g., responsive design) and external open-source libraries based on HTML5 and JavaScript. NEST Desktop 3 is based on the open-source web application framework ‘Vue.js’^5^, which provides a collection of standard GUI widgets and components. The GUI styles offered by ‘Vuetify’^6^ are already used in many other applications and, thus, offer a certain level of consistency in the GUI design for NEST Desktop. The application NEST Desktop runs solely in the web browser of the user. Data of projects, models, and app settings are stored on the local system running the browser.

The visual components of NEST Desktop rely on various JavaScript libraries (Fig. 2C). Graphical representations of neuronal networks use ‘D3.js’^7^. Interactive charts to display simulated activity are realized with ‘Plotly.js’^8^. 3D animated renderings of activity data resulting from simulations of spatially structured networks use ‘Three.js’^9^.

For data handling, NEST Desktop uses ‘PouchDB’ to store data in IndexDB which is by default built into the web browser. ‘PouchDB’ is a JavaScript-based library for ‘CouchDB’ and manages databases with a version-control system.

### Back end implementation

The back end (Fig. 2, yellow rows) hosts first and foremost the simulation engine NEST and it is programmed in Python 3 in conjunction with generic C++. The interface is set up such that the user can directly communicate with NEST via NEST Desktop. NEST Simulator predefines models for neurons, devices, and synapses that are directly selectable in the GUI. Detailed model descriptions can be requested from the NEST Simulator via a RESTful API.

### Development, installation and documentation

The development of NEST Desktop (Fig. 3) follows a community approach. The source code is open-source and available on the GitHub platform^10^ under the MIT License (Fig. 3, middle left). The software development follows the GIT workflow and ‘ESLint’ enforces that style, formatting, and coding standards of the code base are adhered to.

**Figure 3:**
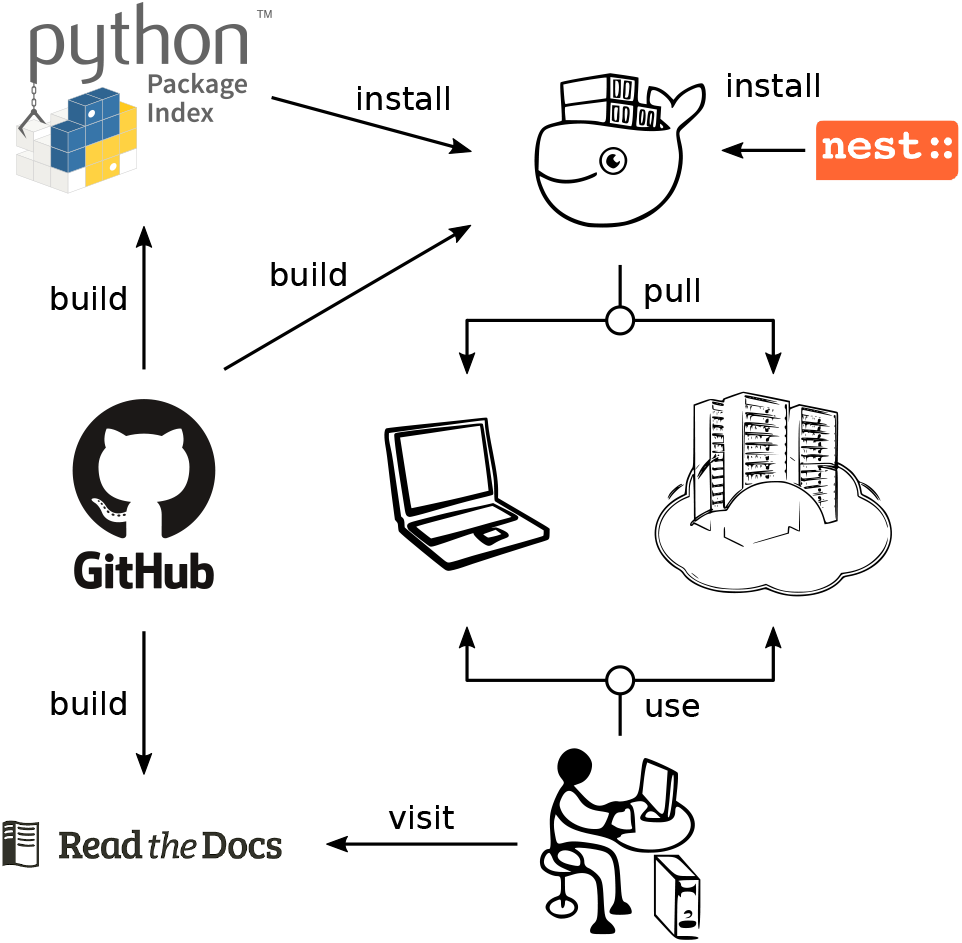
Software development and documentation of NEST Desktop. The code is open-source and available on GitHub. The development of NEST Desktop makes use of Python Package Index and Docker Hub. An associated docker image containing NEST Desktop together with the simulator NEST can be pulled on any local machine or server infrastructure. The ReadTheDocs platform provides detailed information about NEST Desktop for users, lecturers, deployers, and developers.

Running NEST Desktop requires the installation of both the front end NEST Desktop itself and the NEST Simulator as the back end. Both components need to work together. The main reference for installation instructions is the online documentation on the ReadTheDocs platform^11^. Here, we limit ourselves to an overview and highlight some alternative approaches for setting up NEST Desktop. For the easiest local installation, we provide virtual Docker containers (Fig. 3, top right) for NEST Desktop and NEST Simulator which can be installed together using ‘Docker Compose’ with the configuration file ‘docker-compose.yml’ and a single command: `docker-compose up`. Since Docker is available for different operating systems (Linux, Windows, and Mac), this approach allows to provide and use the Linux-based ecosystem of NEST Desktop not only on a local laptop but also on a wide range of other infrastructures (Fig. 3, middle right). Alternatively, NEST Desktop has already been deployed on EBRAINS^12^, the European research infrastructure developed by the Human Brain Project^13^. Everyone with an EBRAINS account can use NEST Desktop there online without any installation. Furthermore, NEST Desktop was temporarily deployed on bwCloud^14^, a university-internal cloud computing resource for teaching purposes. NEST Desktop is installation-free in the sense that a computer center can provide NEST Desktop as a service such that the user only requires a web browser.

As an alternative, NEST Desktop and NEST Simulator can be obtained separately. On Docker Hub, there is a dedicated image for NEST Desktop^15^. NEST Simulator can also be obtained from Docker Hub via the official NEST repository^16^. For advanced users, the front end NEST Desktop is in addition available as a Python Package^17^ published on Python Package Index (PyPI), a third-party software repository for Python, and can be installed with the ‘pip’ package manager (Fig. 3, top left): `pip3 install nest-desktop`. Since NEST 3, a full installation of NEST Simulator on the host system will also provide the API server for RESTful requests. If NEST Desktop and NEST Simulator are installed separately, they can be started with `nest-desktop start` and `nest-server start`, respectively, after which the GUI opens in the web browser and is connected to the simulation engine.

Beyond installation instructions, the documentation of NEST Desktop on ReadTheDocs (Fig. 3, bottom left) explains the usage of NEST Desktop by step-by-step examples using text, animations, and video tutorials. The documentation is organized in separate sections for users, lecturers, deployers, and developers. The user documentation guides users to build networks, parameterize nodes and connections, and perform simulations. Lecturers learn how to deliver course material using NEST Desktop. Deployers find instructions to set up NEST Desktop on a machine via the Python Package or using Docker or Singularity installations instead. Developers get first insights into the code base of NEST Desktop and are welcome to contribute. To facilitate getting started with NEST Desktop, a few example projects with simple network models are also integrated into the tool and can directly be inspected and modified by a new user.

## Results

NEST Desktop implements the whole conceptual workflow of neuronal network simulations (Fig. 1) in a common, graphical user interface running in the web browser (Fig. 4). Users can seamlessly switch between three views with different functionality: the ‘network editor’ (Fig. 4A) for graphical network construction and parameterization, the ‘activity explorer’ (Fig. 4B) for analyzing activity data after a simulation run, and the ‘lab book’ (Fig. 4C) for the project overview. The following provides details on these views and their related functionality, then illustrates a fictive use case about the tool’s employment in the classroom and beyond.

**Figure 4:**
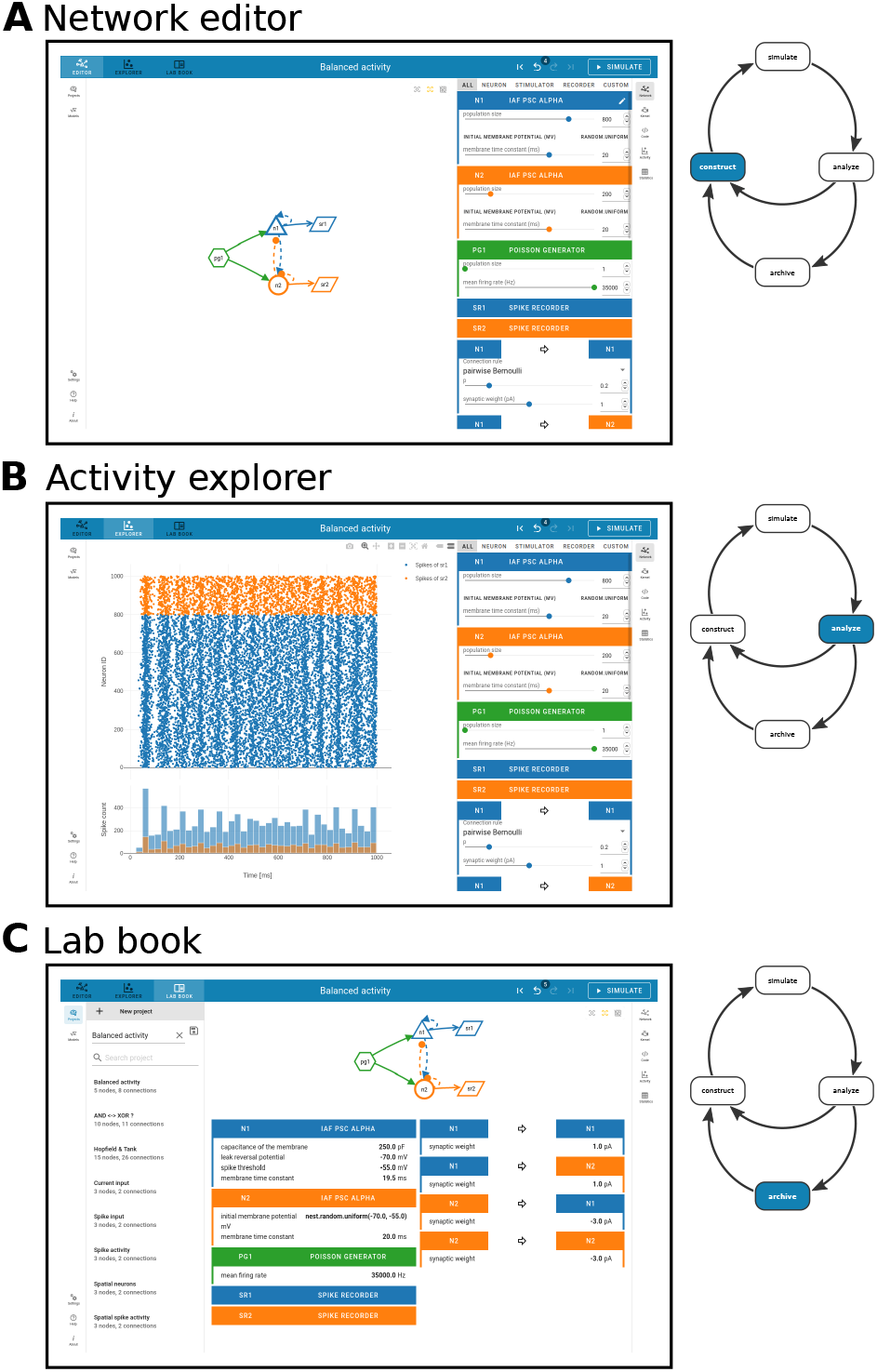
Network editor, activity explorer, and lab book share a common user interface. The steps of NEST Desktop’s conceptual approach for rapid prototyping (right, adapted from Fig. 1) correspond to the distinct views of the tool. **(A)** The network editor provides interactive construction and modification of the neuronal network. **(B)** The activity explorer visualizes network activity. **(C)** The lab book gives users a complete picture of a constructed network. The project manager is shown on the left.

### Graphical construction of neuronal networks in the network editor

The ‘network editor’ allows the visual construction of a network graph by selecting and assembling node and connection elements (Fig. 5A). The appearance of those elements is inspired by the graphical notation proposed by Senk et al. (2020).

**Figure 5:**
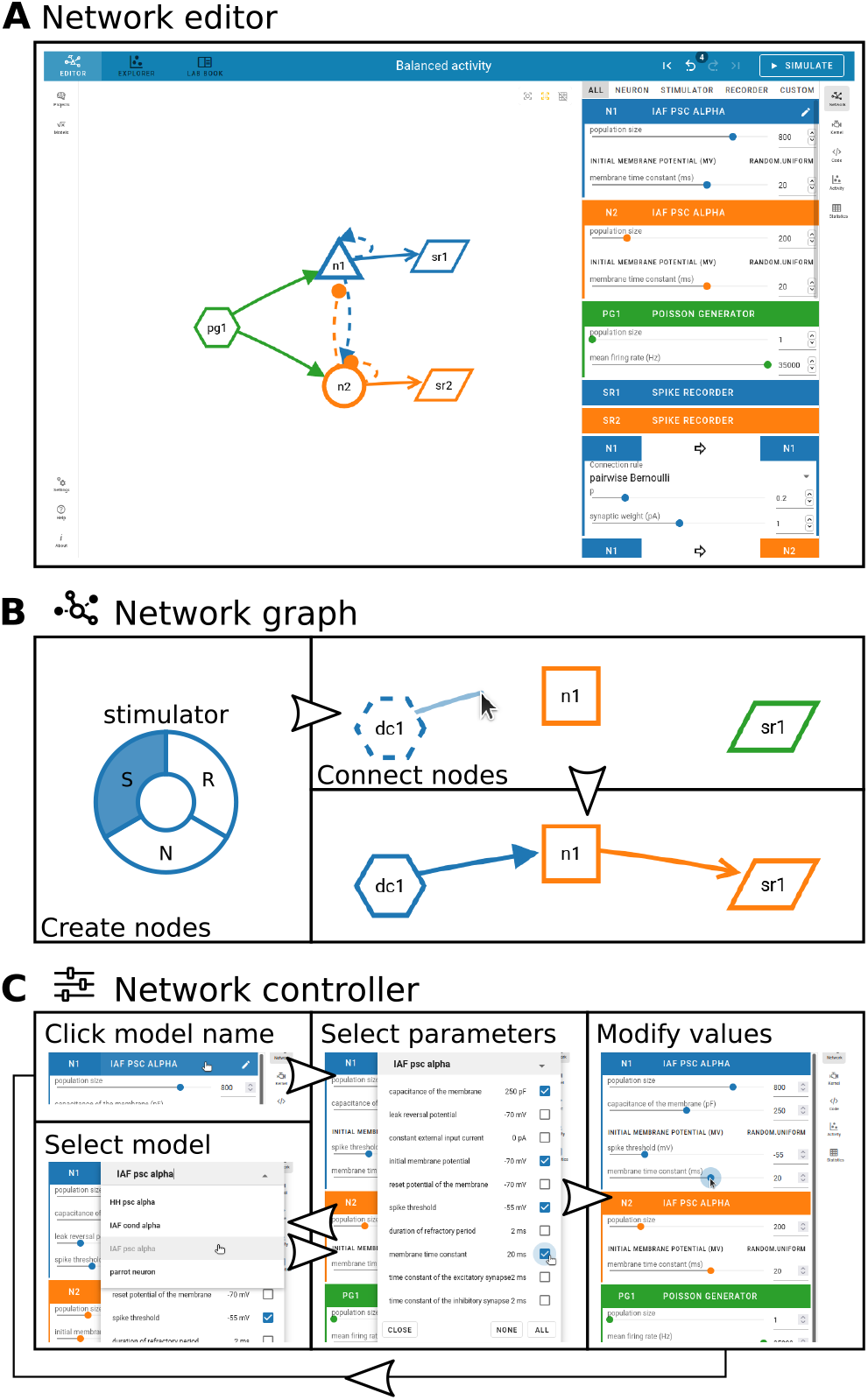
Visual network construction. **(A)** The network editor is the main work space to graphically construct a network graph (left) and adjust network properties with the controller panel (right). Stacks of node and connection parameters associated with the chosen models are displayed in colored panels. **(B)** A right-click with the mouse in the blank space of the editor opens a creation panel (left) to add a node to the network. Node types are stimulator (S), neuron (N), and recorder (R). Connections between nodes are drawn with the mouse cursor (top right). A minimal network may consist of a stimulator, a neuron, and a recorder (bottom right). **(C)** The network controller to the right of the network graph allows users to select and parameterize models. Clicking once on the model name (top left) opens a popup for selecting parameters via checkboxes (middle); clicking there twice allows the user to select a different model from a drop-down menu (bottom left). When a subset of model parameters is selected, the corresponding values can be modified (right) by moving sliders, incrementally increasing and decreasing the value, or by directly entering the value. A restart of NEST Desktop is also possible.

Clicking the right mouse button in the network graph area triggers the creation of a new node (Fig. 5B, left). A pie menu shows the available node types to choose from. Node types are distinguishable by unique shapes. Stimulator (S) nodes have a hexagon shape; they act as input devices that produce signals to be transmitted towards target nodes. Recorder (R) nodes have a parallelogram shape; they represent devices that record signals emitted by other nodes. Neuron (N) nodes integrate the input they receive from other nodes and transform them to recordable outputs. Per default, neuron nodes are of a general type and depicted as squares. The neuron node type can be refined further when nodes are connected. Nodes get distinct colors, which help to associate them with their respective parameter settings and simulated activity throughout the tool.

A directed connection between two nodes is established by clicking first on the connector of a source node and then on a target node (Fig. 5B, right). Selecting the same node both as source and target is also allowed. The arrow representing the connection has the same color as the source node. If all connections from a particular source neuron node to other neuron nodes are parameterized with positive weights, the type of the source neuron node gets refined to be excitatory; if all outgoing weights are negative, the node is inhibitory. Excitatory neuron nodes have triangular shapes resembling pyramidal neurons, and inhibitory ones have a circular shape.

Nodes and connections are configured via the controller panel to the right of the network graph area (Fig. 5A, right). The user can specify properties of the graph elements by choosing predefined models, selecting a parameter subset for these models, and modifying their values (Fig. 5C). One mouse click on the header with the current model name enables the parameter selection for that model, and a second click opens a menu for changing the model. The available models depend on the node type (stimulator, neuron, or recorder), and each of them has its own set of parameters. The models are all part of NEST, and the user can query the available model descriptions from the NEST source code. A neuron node, for instance, may represent a whole population of individual neurons sharing the same model. The parameters can then either be the same for all neurons of the population or sampled from an array or from a random distribution (in “expert mode”). Optionally, users can also assign spatial positions to neurons or stimulating devices in the network.

The user can specify properties of the graph elements by, first, choosing predefined models via a drop-down menu and, second, adjusting its parameter values with sliders or by directly typing the numbers (Fig. 5C). A neuron node, for instance, may represent a whole population of individual neurons sharing the same model. Each of the available models is part of NEST and has its own set of parameters and the user can query the available model descriptions. Optionally, users can assign spatial positions to neurons or stimulating devices in the network.

During editing, each change of the network is logged such that the user can go back in history and undo and redo changes.

### Code generation from the network graph

The network graph is automatically rendered into executable PyNEST (Eppler et al., 2009) code with a direct correspondence between graphical elements and textual code snippets (Fig. 6). PyNEST is the Python interface to the simulation engine NEST (Gewaltig and Diesmann, 2007). The script is structured in blocks that are produced in the same order as they are executed in the back end when a simulation run is inquired. First, the technical setup of the simulator is added to the script: modules are imported and further parameters can be defined to be passed to the simulation kernel. The following code lines in Fig. 6 account for creating and connecting the nodes as defined by the user in the network editor. Afterwards, the command for initiating the state-propagation phase is defined, which is the actual simulation. The last block contains the code for collecting the recorded network activity data for subsequent visualization and analysis in the ‘activity explorer’. Clicking the ‘Simulate’ button triggers the execution of this code with NEST.

**Figure 6:**
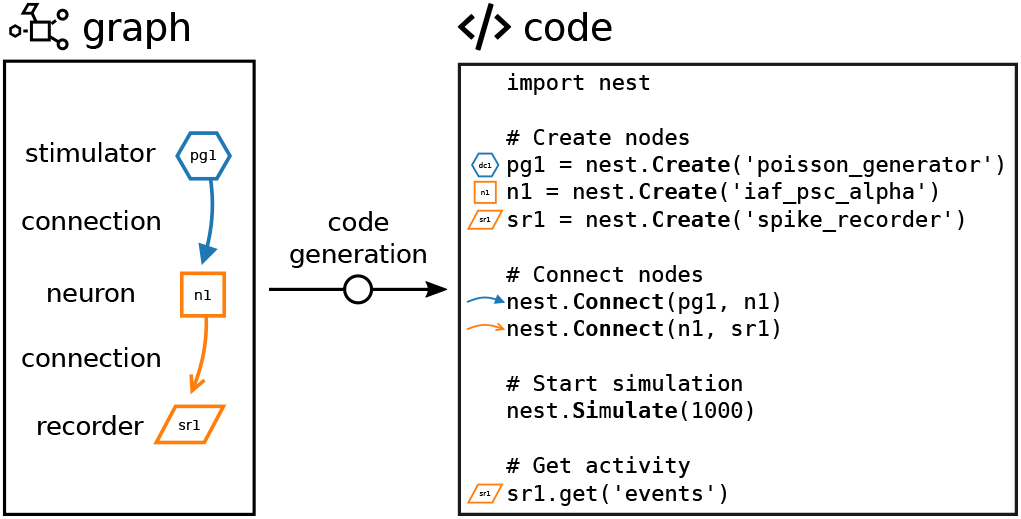
Code generation through visual network construction. The graphically composed network (see Fig. 5) is automatically translated into textual source code. Visual elements in the network graph (shapes for nodes and arrows for connections) are associated with generated code lines. The resulting script is a complete definition of a simulation experiment in PyNEST with code blocks to be executed in succession: ‘Create nodes’, ‘Connect nodes’, ‘Start simulation’, and ‘Get activity’. The sketched network of only three connected nodes (stimulator to neuron to recorder) is a minimal example for illustration; further details such as parameter values set via the GUI are also turned into code.

### Interactive data analysis with the activity explorer

Dependent on properties and parameterization of recording devices in the constructed network, different types of activity data are returned from NEST for inspection in the activity explorer. The data comprises unique IDs of sending neurons and so-called events, which are either spikes (discrete time stamps) or quasi-analog signals, e.g., membrane potentials (sampled in given time intervals). The charts in the bottom left panel of Fig. 7 show vertically arranged traces of membrane potentials as line graphs, a spike raster as scatter plot, and computed spike counts across time as histogram. If the data additionally contains neuron positions in 2D or 3D space, the activity can also be animated in a 3D graph (Fig. 7, bottom right). Beside the visual analysis, NEST Desktop also has the possibility to display basic spike-train statistics in table format. The top right panel of Fig. 7 demonstrates such a table with statistics calculated from the raw data.

**Figure 7:**
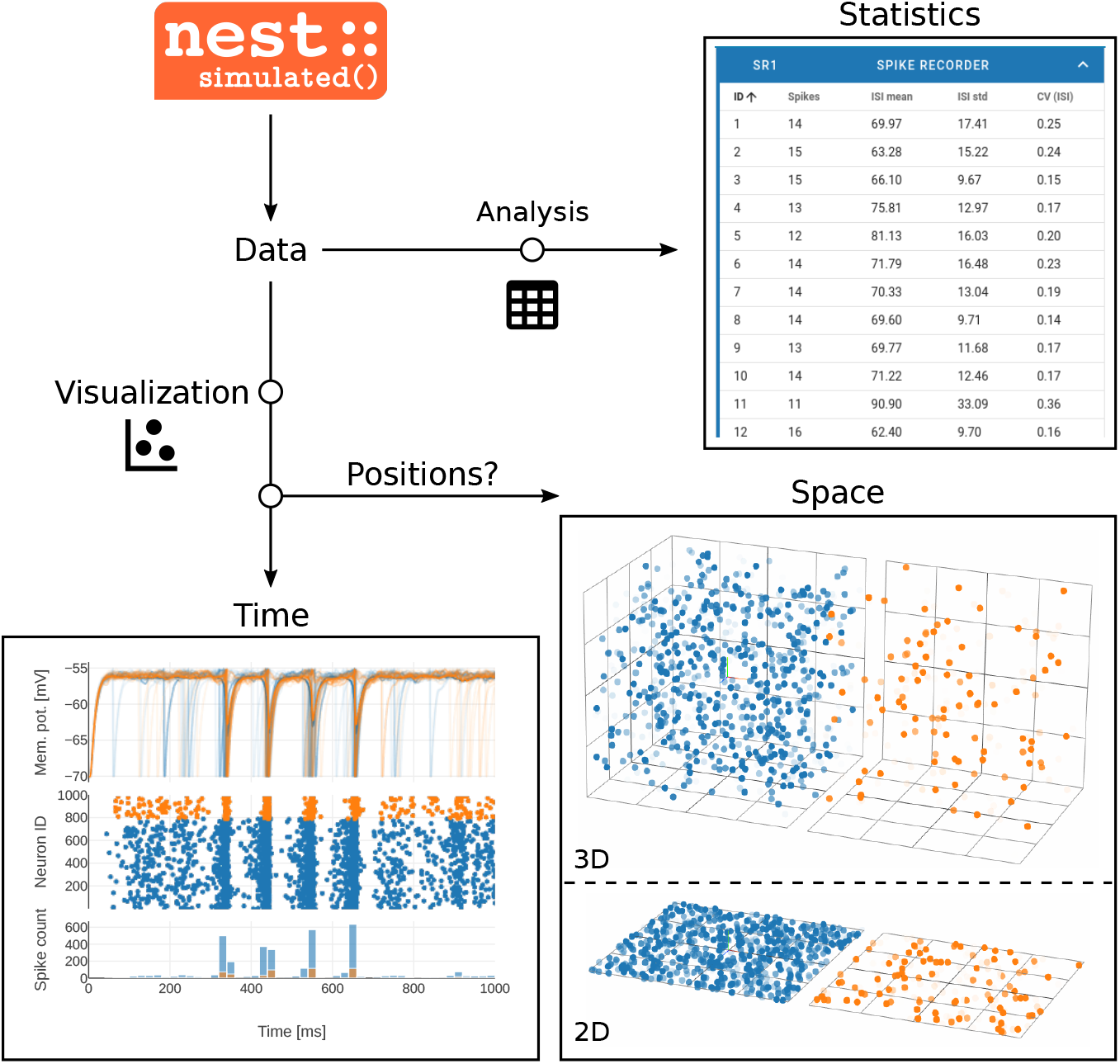
Data analysis and visualization. The NEST simulator executes the network simulation and returns recorded activity data to be analyzed and visualized (top left). Quasi-analog signals like membrane potentials and discrete spike times can be displayed across time (bottom left). Such visualization is accompanied by basic analysis like the computation of spike counts. If the neurons in the simulated network are arranged in space, a 2D or 3D animation offers a view of the ongoing activity at the respective neuronal positions (bottom right). Calculated quantities are presented in table format (top right).

### Project management and image export

NEST Desktop sessions are treated as projects and handled by the project manager (Fig. 8A, top): one can either start a new project and construct a network from scratch or load a previously saved project to extend an existing network structure. Existing projects, set up and saved at another time or on another machine, can be duplicated, updated, or deleted. Projects are synchronized with a built-in database in the browser on the client-side, but they can also be exported to and loaded from file (Fig. 8A, bottom).

**Figure 8:**
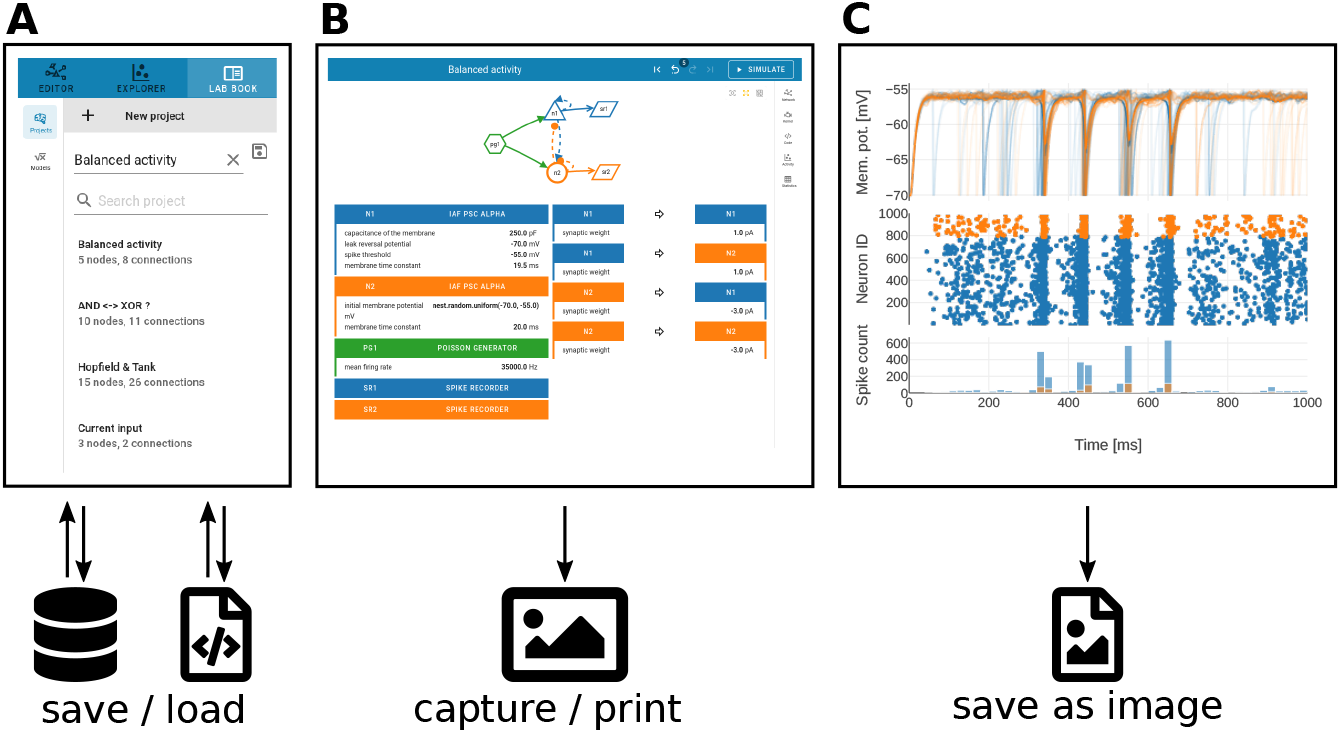
Project archiving and image export. **(A)** Previously constructed networks (see Fig. 5) can be stored in a database within NEST Desktop or exported to a file for later reloading. An example list of saved, loadable projects is shown. **(B)** The network graph and its description can be captured as screenshot. **(C)** Charts visualizing activity data (see Fig. 7) allow for export to either a rendered image (‘.png’) or a vector graphic (‘.svg’).

Apart from saving the status of a project, NEST Desktop also encourages the export of figures showing network definitions or activity charts to protocol observations. Particularly suitable for this purpose is a view that resembles a ‘lab book’ (Fig. 8B): the graphical representation of the network is here displayed above a two-column table specifying nodes and connections to provide a compact overview. For capturing the graph and its parameters, we recommend using an external screenshot tool or printing to file from the browser (Fig. 8B). For saving activity charts, however, NEST Desktop provides internal functionality: those figures can be exported directly as high-quality vector-graphics or as pixelated images (Fig. 8C).

### Use case: NEST Desktop in the classroom

Here, we illustrate how NEST Desktop may be employed as a learning and teaching tool in a hands-on session of an introductory course of computational neuroscience. The students are expected to have only limited prior knowledge in the field and the lessons are supposed to teach them the principles of spiking neuronal networks. Computer simulations are intended to help them develop an intuitive understanding of the network dynamics. The lesson discussed here aims to construct a network of two interconnected populations of leaky-integrate-and-fire (LIF) neurons driven by an external input. The activities of the excitatory and inhibitory neuron populations should be balanced. This scenario describes a classical example of an emergent property in a rather simple network configuration (Vreeswijk and Sompolinsky, 1996; Brunel, 2000). Our fictional student Noel is highly motivated to learn about this topic and the method of neuronal network simulations, but he is inexperienced in programming. We will explain how NEST Desktop helps Noel to achieve the goal nevertheless.

The course takes place in the university’s computer lab and has been prepared by the tutor Juno. She consulted the documentation of how to deploy NEST Desktop in a virtual machine on computer resources provided for students (Fig. 3, ReadTheDocs) and found a pre-built container with the tool (Fig. 3, whale, Docker). After following a few steps, NEST Desktop is ready to be used by the students without the need of manual installation or configuration (Fig. 3, laptop/cloud).

Noel opens the network editor (Fig. 5A) and begins to set up the network. In the two-dimensional scratch panel, he creates one neuron node and one recording device per population to track the neuronal activity (Fig. 5B, left). He adds a single stimulus device to provide external input to both populations. The next step is to connect the neuron and device nodes (Fig. 5B, right). The connectivity between network nodes can be defined with different deterministic and probabilistic rules selectable from a drop-down menu. Each neuron node is connected to the other one and to itself; the neurons are connected randomly (pairwise Bernoulli) with a given connection probability. Noel notes that the nodes are differently labelled and colored which helps matching nodes and connections with the information shown in the other panels (Fig. 5).

Subsequently, Noel opens the network controller and specifies the models represented by the nodes. He finds the neuron model he is looking for: ‘IAF psc alpha’, a current-based ‘leaky integrate-and-fire’ neuron with alpha-shaped post-synaptic currents (Fig. 5C, left bottom). As an alternative to this LIF neuron model, a Hodgkin-Huxley neuron model is also available, which has more biophysical details. Noel chooses model parameters which are relevant for the exercise (Fig. 5C, middle). These selected parameters can then be modified from their preset default values by either using sliders, or by typing the intended values into the input field (Fig. 5C, right).

An important parameter is the number of elements in a node, which is also referred to as population size. In this example, the excitatory population is larger than the inhibitory one, but inhibitory connections are stronger for compensation. Noel sets the population sizes of both neuron nodes accordingly and also modifies other parameter values where necessary.

In the code editor, Noel finds the scripting code that is automatically generated from the graphically constructed network (Fig. 6). Every visual element has its respective counterpart in the script and Noel recognizes the model names and the parameters he has set earlier via the GUI. Noel finds out that he can modify the code in the editor directly. Just for testing this option, he changes the value of one parameter. Noel learns that the programming language of this script is called PyNEST. As the network is now completely defined, Noel clicks the ‘Simulate’ button, which triggers the transmission and the execution of the PyNEST script in the background.

After the simulation, which only took a few seconds, Noel starts exploring the recorded network dynamics in the activity explorer. During network construction prior to the simulation, he has chosen spike recording devices and focuses now on analyzing the spiking activity. A raster plot shows the spike times of both neuronal populations (Fig. 7, bottom left). In this plot, Noel registers noise-like activity of both neuronal populations. He pans the plot window to the episode of interest, zooms in on some individual neurons in the network and observes that they emit spikes at non-synchronous, seemingly random time points. The subjacent histogram displays the spike count in time bins, accounting for all neurons in each population. Noel interactively changes the bin width and observes how the spike count adjusts. Although individual neurons only occasionally contribute a spike, the population spike counts are stationary over time reflecting the balance between excitation and inhibition. Via the side navigation, Noel opens a table showing statistical measures of the spike data of individual neurons (Fig. 7, top right). The coefficient of variation of the inter-spike intervals (CV_ISI_) is just below one for most neurons, indicating that the spiking activity in the network is almost as irregular as a Poisson process (Softky and Koch, 1993). Noel concludes that both neuronal populations operate in the asynchronous-irregular (AI) dynamical regime (Brunel, 2000). Next, Noel returns to the network editor, adds a stimulus device to apply negative currents to the inhibitory population during a defined period of time. He also adds multi-purpose measurement devices (‘multimeter’) to record the neurons’ membrane potentials as quasi-analog signals. He re-simulates and observes changes in the network activity: during the stimulation phase, both populations exhibit highly synchronous and oscillatory behavior, visible in the membrane potential traces, the spike raster, and the population spike count histogram (Fig. 7, bottom left). As a last test, he randomly assigns positions to the neurons and observes the animated activity resolved in space (Fig. 7, bottom right).

Noel finishes the exploration and analysis of network dynamics and saves the project with a descriptive name (Fig. 8A). The project management panel allows him to reload the project later to resume the exploration at the point where it was stopped. He also exports the project to a file in order to load it on his home computer later, or to share it with another student. Noel’s final task is to document his exploration of the balanced network model and he writes a report about the simulation setup and the analysis of the simulated network activity. To enhance the protocol with graphics, Noel first uses the built-in screenshot option to capture the lab book as an overview of the network (Fig. 8B). A display of the simulated data should have high quality to resolve important details of the spiking activity. Noel finds that he can export the charts as Scalable Vector Graphics (.svg) which meets that requirement (Fig. 8C). Ultimately, Noel includes the figures into his protocol and moves on to the next lesson.

### NEST Desktop beyond teaching

Here, we provide a short outlook on the potential usage of NEST Desktop beyond its major teaching purpose. Juno is a researcher and, apart from teaching courses, she studies spiking network models for her own scientific work. NEST Desktop has proven useful more than once in quickly investigating certain features of the simulator NEST, or testing the dynamics of a toy network before integrating the insights into larger models expressed in scripted code. If she is not familiar with the correct NEST syntax, she has even found herself constructing the respective network parts graphically in NEST Desktop and obtaining the executable PyNEST code from the build-in code-generation functionality. But NEST Desktop does not only help Juno to acquire a better intuition for her models, she also uses the tool for explaining her work to others. She finds that the audience can better grasp network structures and mechanisms behind activity dynamics if presented interactively. After her talks, she shares network configuration files with interested members of the audience who can then continue exploring the shown networks with NEST Desktop on their own machines. Thus, NEST Desktop can also support the daily routine of researchers in various aspects.

## Discussion

NEST Desktop is an interactive, web-based Graphical User Interface (GUI) to the neuronal network simulation code NEST, primarily developed for teaching the fundamentals of computational neuroscience. Students can choose from a number of available neuron, device and synapse models, combine them into network structures, and set custom parameters. The graphically constructed network is automatically converted into scripted code which the simulation engine NEST executes in the background. Simulation results are returned to the GUI, where the students can explore neuronal activity with a selection of analysis tools. Hence, our approach demonstrates a conceptual marriage of a powerful simulation engine and an intuitive, user-friendly GUI.

The use case “NEST Desktop in the classroom” which is described in the Results section is based on the actual use of NEST Desktop for teaching computational neuroscience as part of the university education of bachelor and master students and in independent tutorials. Particular challenges of these courses are the very heterogeneous levels of programming skills and background knowledge in neuroscience among the participants. NEST Desktop has already proven to support teaching successfully both in the classical classroom setting with physical attendance and in online formats. Online formats have been boosted due to the COVID-19 pandemic and NEST Desktop has shown itself to be a valuable tool in this situation. In online courses, the students have the chance to contact the tutors and lecturers using video conference tools or via a messenger channel to get answers to their questions and discuss problems regarding the course content or NEST Desktop usage. All these teaching events put NEST Desktop to the test. Gathering feedback from students helps to identify shortcomings and drives the development of the tool. We have observed that the students generally show a reasonably fast learning success making a one-week course feasible. They are typically able to operate NEST Desktop independently already on the first day. The experience they gain from exploring networks using the tool helps them to answer questions from computational neuroscience posed to them in the course script. In an informal round of feedback, students attested NEST Desktop a good level of usability and they gave various positive comments on their user experience. However, there is still room for improvement due to the limited feature set of NEST Desktop as exposed by the students’ feedback.

Here, we contrast NEST Desktop to the standalone application NEST Simulator. NEST Desktop builds on the PyNEST interface of NEST Simulator and can therefore provide access to most of its functionality. The translation of Python commands into elements of the GUI includes manual steps for the developers of NEST Desktop. For reasons of clarity and comprehensibility, not the whole multitude of neuron and synapse models and lower level commands available in NEST Simulator have a GUI counterpart, but only a representative subset that can be extended if needed. Multi-compartment neuron models and synaptic plasticity, for example, are currently not accessible. The set of models in NEST Simulator itself can be extended with NESTML (Plotnikov et al., 2016).

Furthermore, each simulation experiment defined in NEST Desktop is self-contained and comprises all steps (network construction, simulation phase, and retrieval of activity data) of a digitized scientific workflow. Plain NEST is in that sense more flexible, as a running simulation can be interrupted to change parameters and resumed if desired. The PyNEST code can also be combined with generic Python code in case that a required functionality is not yet available in NEST but can be achieved by combining low-level commands of the PyNEST API.

Besides, the size and complexity of networks which can be simulated with NEST Desktop are limited by the hardware resources accessible to the NEST Simulator back end; typically these resources are laptop-equivalent or correspond to one compute node. While NEST Simulator lends itself to simulations of large networks with millions of neurons using high-performance compute clusters and parallelization with MPI and OpenMP, NEST Desktop currently only supports pure multi-threading for NEST Simulator. The attempt to simulate too large networks leads to inconveniently long simulation times and eventually even exhausts main memory. On that account, the GUI provides reasonable default ranges for population sizes. Although generally valid numbers cannot be given, we can conservatively state that networks on the order of a few thousand neurons can routinely be simulated with NEST Desktop. The visualization performance of the network activities is also limited by data size.

Although theoretically not forbidden in NEST Desktop, it may become impractical to construct complex networks in the GUI that consist of a large number of distinct and differently parameterized neuron populations. To address these problems and alleviate the procedures, NEST Desktop provides the possibility to clone nodes during network construction and to customize which nodes and connections are shown for setting parameters.

Regarding data analysis, both NEST Desktop and NEST Simulator provide only basic plotting routines to check simulation results. Given its interactivity and simple statistical analysis, the GUI provides more features than plain NEST. For reasons of modularity, detailed analyses are outsourced to separate, specific tools. NEST Desktop has been designed for learning the fundamentals of simulation and for small proof-of-concept simulation studies. In this spirit, NEST Desktop facilitates the daily routine of a researcher. However, for advanced simulations of large networks, full access to all features of the simulation engine and more flexibility may be required; here the script-based approach of NEST Simulator is recommended.

Based on the useful feedback from given courses and beyond, we identify the following concrete directions in which the development of NEST Desktop may continue: while NEST Desktop already strictly separates the GUI from the simulation engine, one could even expand on the modularity. A possible next step would be to separate GUI and data analysis, as well as storage. The front end engine obviously has limited capability for advanced mathematical operations, like computing spike train correlations or the power spectrum using the Fast Fourier Transform (FFT). An interface to the Python-based data analysis community toolbox Elephant (Denker et al., 2018), which offers such functionality, therefore seems to be a more appropriate and general solution.

In computational neuroscience, several GUIs have already been developed over the last two decades, mostly linked to a specific simulation engine (see Table 1). Most modern GUIs run in a web-browser (HTML) and are therefore platform-independent and installation-free. Table 2 identifies which features of NEST Desktop are available to some extent also in the other tools existing. The design focus, i.e., whether they target rather the visualization of the network graph or the activity, is different between these tools, although many show some functionality of both. The earlier graphical interface ‘SLIDE’ (Matyak, 1996; Gewaltig et al., 1996) for NEST has not been developed further but next to network structure and activity introduced a third aspect: the protocol of a virtual experiment. These ideas were inspired by visual programming at the time. To our knowledge they have not been picked-up again in the context of neuronal network simulation, but movie editing software like ‘Blender’ (Bruns, 2020, the similarity was pointed out by Marc-Oliver Gewaltig in private communication) has aspects of this. Because of the problems in stabilizing and maintaining graphical user interfaces in the middle of the 1990s (Senk et al., 2018, contains some review) NEST development has primarily focused on the independent simulation engine. ‘Open Source Brain’ (Gleeson et al., 2019), ‘Neuron UI’ and ‘NetPyNE UI’ (Dura-Bernal et al., 2019) are extensions of ‘Geppetto’ (Cantarelli et al., 2018) framework but ‘Neuron UI’ appears to be no longer in development. From the user perspective, the tools ‘Open Source Brain’, ‘NetPyNE UI’ and ‘Nengo GUI’ (Bekolay et al., 2014) follow a similar approach as NEST Desktop. NEST Desktop, however, is unique in that from the perspective of the user it is installation-free if deployed on a server infrastructure. The user only requires a browser and has access to advanced compute resources independent of local capabilities. This software architecture makes NEST Desktop well suited for a classroom setting. Furthermore, the GUI enables the user to directly access the script showing a one-to-one correspondence between graphical elements and textual code snippets. This not only enables additional modification to the generated code before sending to the back end (as described in the use case) but also gives the learner the opportunity to gather first-hand experience with the actual code. This facilitates the later step to programming PyNEST scripts without the need of NEST Desktop and, thus, enables the creation and simulation of highly complex models with NEST as is the case in most scientific use cases of NEST. Finally, in contrast to the other outlined projects, the primary motivation of our work is to create a self-contained educational tool using the language and symbols of the problem domain.

**Table 2:**
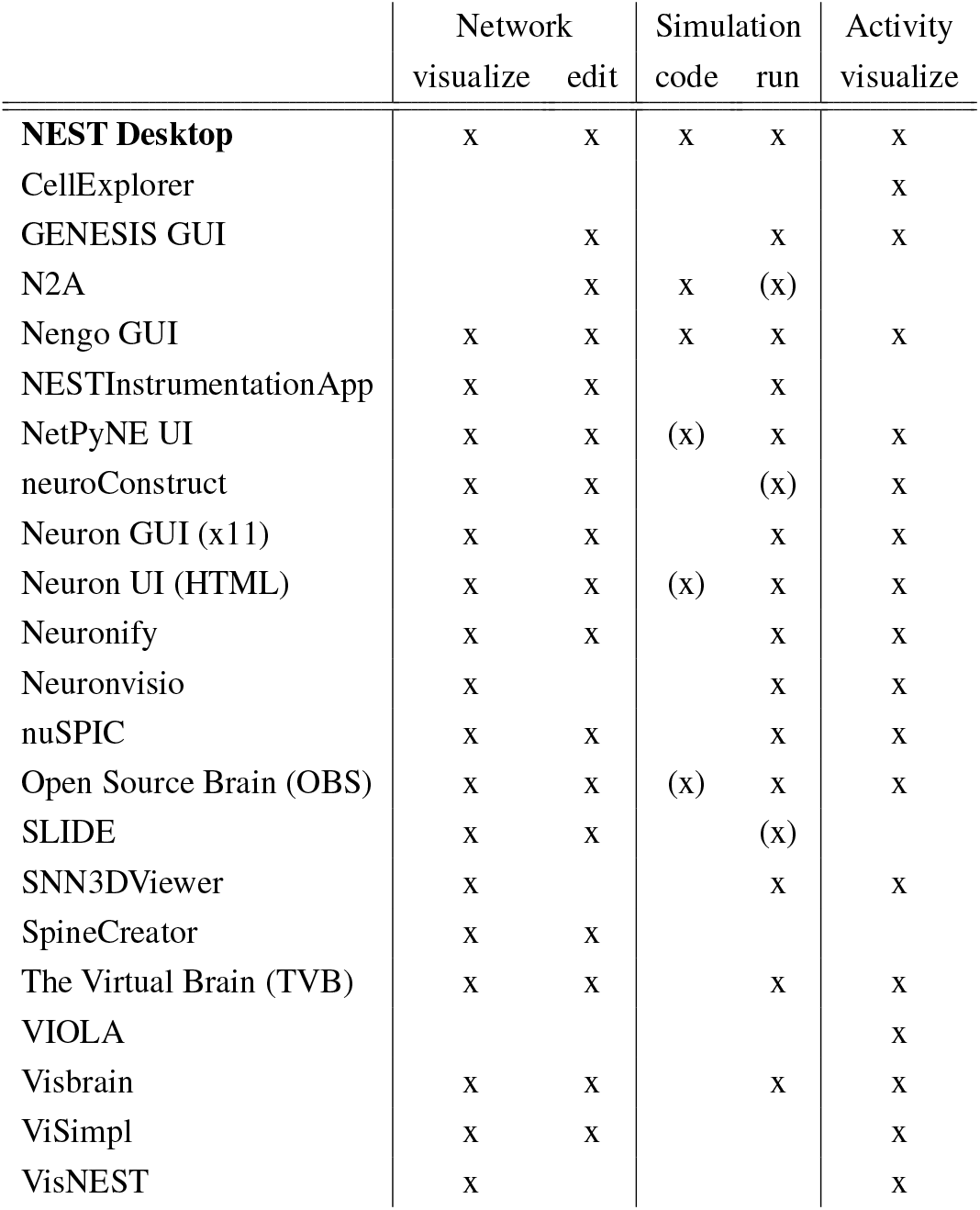
Characterization of GUIs. The GUIs from Table 1 (here sorted alphabetically after NEST Desktop) are compared based on which steps of network construction, simulation, and activity analysis they cover. The network aspect is split up into visualizing networks and the possibility to edit them by means of the GUI. For the simulation step, the table distinguishes between a feature to generate and display simulation code and the option to actually run a simulation. The marker ‘(x)’ in the simulation column means that (executable) code is provided but rather debug code or console instead of the actual simulation code.

NEST Desktop supports teaching of computational neuroscience by making computer simulations intuitively accessible. The use of NEST Desktop actually reverses the sequence of skills to be learned. Courses can now directly start with scientific content, without students having to learn scientific programming first. Once the students have developed their intuition for neuronal network models, it is much easier for them to get started with the actual scripting of simulations and conduct more sophisticated research projects in the field on their own.

## Acknowledgments

We thank Jens Buchertseifer for the collaboration in code development and review of NEST Desktop, and Jochen Martin Eppler for discussion and development of NEST Server. Moreover, we thank Sara Konradi, Jessica Mitchell, Dennis Terhorst, Steffen Graber and the community of NEST developers for discussion, review of the user documentation and the web page on EBRAINS. At Freiburg, we thank tutors and students of the Bernstein Center for beta-testing NEST Desktop, enduring the early days, and valuable feedback. We further gratefully acknowledge the service of the HBP Support Team for deployment on EBRAINS as well as the operators at the Rechenzentrum of University of Freiburg for the deployment on statewide bwCloud infrastructure. Finally, we thank Thomas Matyak and his aunt for uncovering the thesis on SLIDE, and the research assistant (HiWi) Peter Bouss for finding Thomas.

## Funding sources

This project has received funding from the European Union’s Horizon 2020 Framework Programme for Research and Innovation under Specific Grant Agreement No. 785907 (Human Brain Project SGA2) and No. 945539 (Human Brain Project SGA3). This project was funded by the Helmholtz Association Initiative and Networking Fund under project number SO-092 (Advanced Computing Architectures, ACA). This work was supported by the DFG Excellence Cluster BrainLinks-BrainTools (grant EXC 1086). The HPC facilities are funded by the state of Baden-Württemberg through bwHPC and DFG grant INST 39/963-1 FUGG.

## Footnotes

1 https://github.com/nengo/nengo-gui

2 https://github.com/nest-desktop/nest-desktop

3 https://github.com/compneuronmbu/NESTInstrumentationApp

4 https://github.com/MetaCell/NEURON-UI

5 https://vuejs.org

6 https://vuetifyjs.com

7 https://d3js.org

8 https://plot.ly/javascript

9 https://threejs.org

10 https://github.com/nest-desktop/nest-desktop

11 https://nest-desktop.readthedocs.io

12 https://ebrains.eu/service/nest-desktop

13 https://humanbrainproject.eu

14 https://www.bw-cloud.org

15 https://hub.docker.com/r/nestdesktop/app

16 https://hub.docker.com/r/nestsim/nest

17 https://pypi.org/project/nest-desktop

## Notes

https://nest-desktop.readthedocs.io

### Competing Interest Statement

The authors have declared no competing interest.

### Summary of Updates

Revision from Reviewer's reply.

